# Mammalian kinetochores count attached microtubules in a sensitive and switch-like manner to control cell cycle progression

**DOI:** 10.1101/463471

**Authors:** Jonathan Kuhn, Sophie Dumont

**Affiliations:** Tetrad Graduate Program, University of California, San Francisco, San Francisco, CA, 94143; Department of Cell and Tissue Biology, University of California, San Francisco, San Francisco, CA, 94143; Department of Cell and Molecular Pharmacology, University of California, San Francisco, San Francisco, CA, 94143

## Abstract

The Spindle Assembly Checkpoint (SAC) maintains genomic integrity by preventing anaphase until all kinetochores attach to the spindle. What specific signals are required for SAC satisfaction at mammalian kinetochores, and in what magnitude, are not well understood and central to understanding SAC signal processing and function. Here, we directly and independently tune candidate input signals – spindle forces and Hec1-microtubule binding – and map SAC outputs. By detaching microtubules from the spindle, we first demonstrate that the SAC does not respond to changes in spindle pulling forces. We then tune and fix the fraction of Hec1 molecules capable of microtubule binding, and interpret SAC output changes as coming from changes in binding, and not spindle forces. While the speed of satisfaction reduces with fewer attached microtubules, the kinetochore turns off the SAC even with few – approximately four – such microtubules. Thus, the mammalian kinetochore responds specifically to microtubule binding, and does so as a single, switch-like, sensitive unit. This may allow the kinetochore to rapidly react to attachments and maintain a robust response despite dynamic microtubule numbers.

## Introduction

The Spindle Assembly Checkpoint (SAC) protects genomic integrity by preventing anaphase until all kinetochores attach to the spindle (Rieder et al., 1995). Unattached kinetochores, and certain improperly attached ones, recruit Mad1 and Mad2, which generate an anaphase-inhibitory signal (Chen et al., 1998; De Antoni et al., 2005; Maldonado and Kapoor, 2011). By metaphase, mammalian kinetochores bind 15-25 microtubules (McEwen et al., 1997; Wendell et al., 1993), are under spindle forces, and have lost Mad1/2. What specific signals are required for SAC satisfaction, and in what magnitude, are not well understood and are key to understanding SAC signal processing and function. One challenge to defining how a given input signal quantitatively maps to SAC outputs is that of directly controlling and tuning candidate signals – spindle forces and microtubule plus-end binding (London and Biggins, 2014) – inside cells. Another is that these candidate signals are interdependent (Akiyoshi et al., 2010; King and Nicklas, 2000) and hard to uncouple.

There are many indications that force across sister kinetochores (across the centromere) is not required for SAC satisfaction (Etemad et al., 2015; Magidson et al., 2016; O’Connell et al., 2008; Tauchman et al., 2015; Waters et al., 1998), and that spindle force on a kinetochore is insufficient for SAC satisfaction without proper attachment (Kuhn and Dumont, 2017). However, whether force from the spindle on a single kinetochore is necessary for SAC satisfaction in normal conditions remains unclear. For example, while mono-attached kinetochores can satisfy the SAC despite not generating force across the centromere (Etemad et al., 2015; O’Connell et al., 2008; Tauchman et al., 2015), this could be caused by opposing forces on *individual* kinetochores through outward spindle forces on chromosome arms (Cane et al., 2013; Drpic et al., 2015; Maresca and Salmon, 2009; Uchida et al., 2009). Further, inhibiting pulling forces by suppressing microtubule dynamics does not prevent SAC satisfaction (Magidson et al., 2016; Waters et al., 1998), but this could be due to altered dynamics. Finally, recent evidence suggests that centromere tension may still play a role in certain scenarios (Janssen et al., 2018). Better spatial and temporal control of microtubule pulling forces on kinetochores – without affecting microtubule binding – is required to determine whether spindle forces are required for SAC satisfaction.

Regardless of the spindle force generated, kinetochore-microtubule interactions are essential for controlling Mad1/2 levels (Etemad et al., 2015; Kuhn and Dumont, 2017; Tauchman et al., 2015; Waters et al., 1998). The kinetochore protein Hec1/NDC80 is key both for microtubule plus-end binding (Cheeseman et al., 2006; DeLuca et al., 2006) and SAC signaling (McCleland et al., 2003). How Hec1-microtubule interactions cooperate to form a kinetochore-fiber (k-fiber; microtubule bundles attaching to chromosomes) and pull on a kinetochore, and how they inform SAC signaling output and kinetics, are poorly understood. In mammalian cells, the number of kinetochore-microtubules (input) is widely variable (analog), unlike in budding yeast where it is zero or one (Peterson and Ris, 1976): each metaphase kinetochore binds ~20 microtubules (McEwen et al., 1997; Wendell et al., 1993), with around 80 of its ~250 Hec1 copies binding microtubules at a given time (Suzuki et al., 2015; Yoo et al., 2018). The number of kinetochore-bound Mad1 molecules and the cytoplasmic SAC strength are also analog (Collin et al., 2013; Dick and Gerlich, 2013; Heinrich et al., 2013). We know that satisfaction can begin with half of a full microtubule complement (Dudka et al., 2018; Kuhn and Dumont, 2017), but do not know the quantitative relationship between microtubule input and SAC output. To map this relationship, it is necessary to fix and control the normally dynamic number of kinetochore-microtubules without perturbing microtubule dynamics or essential kinetochore functions.

Here, we develop approaches to acutely remove spindle forces on individual kinetochores while maintaining microtubule attachment, and to control the level of Hec1-microtubule occupancy – while quantifying the SAC’s output. We demonstrate that spindle force generation is not required to maintain SAC satisfaction, suggesting that Mad1 levels are only linked to microtubule attachment. By tuning and fixing the quantity and binding affinity of Hec1 at individual kinetochores, we gradually adjust the number of attached microtubules: we show that while reducing this number slows Mad1 loss rates, only four or fewer attached microtubules are ultimately required to satisfy the SAC. Thus, the mammalian kinetochore acts as a single processing unit that responds exclusively and sensitively to microtubule attachments in a manner that ensures accuracy while preventing mitotic delays.

## Results

To determine whether the SAC responds to spindle pulling forces on kinetochores, we developed a laser ablation assay to mechanically isolate a k-fiber from a metaphase spindle in mammalian PtK2 cells (Fig. 1A, Video 1). We expressed EGFP-tubulin and Hec1-EGFP (in a Hec1 RNAi background (Guimaraes et al., 2008)) to assist k-fiber ablation and response tracking. Based on previous work (Elting et al., 2017; Kajtez et al., 2016), we cut the k-fiber close to its kinetochore to minimize residual spindle connections and forces; without these the ablated k-fiber (stub) cannot exert pulling forces to move kinetochores. Unlike previous approaches, removing pulling forces prevents force generation on individual mature metaphase kinetochores regardless of pushing forces on chromosome arms (Rieder et al., 1986). If SAC satisfaction requires spindle pulling forces, acutely removing them on attached kinetochores should re-recruit Mad1/2. To allow time for Mad1/2 recruitment (^~^1.5 min) (Dick and Gerlich, 2013; Pines and Clute, 1999), we prevented force generation from k-fiber-spindle reincorporation (Elting et al., 2014; Sikirzhytski et al., 2014) by repeatedly ablating the k-fiber and partially knocking down NuMA (Elting et al., 2014), a protein essential for reincorporation (Hueschen et al., 2017). As predicted for the removal of spindle connections, during ablation the interkinetochore (K-K) distance dropped to values comparable to that in nocodazole (1.11±0.04 μm) (Fig. 1A-B), the disconnected kinetochore persistently moved away from its pole (Fig. 1C) (Khodjakov and Rieder, 1996), and we could not detect significant microtubule intensity between the k-fiber stub and spindle body (Fig. 1A, Video 1). Thus, the above method removes productive force generation at individual kinetochores.

**Figure 1:**
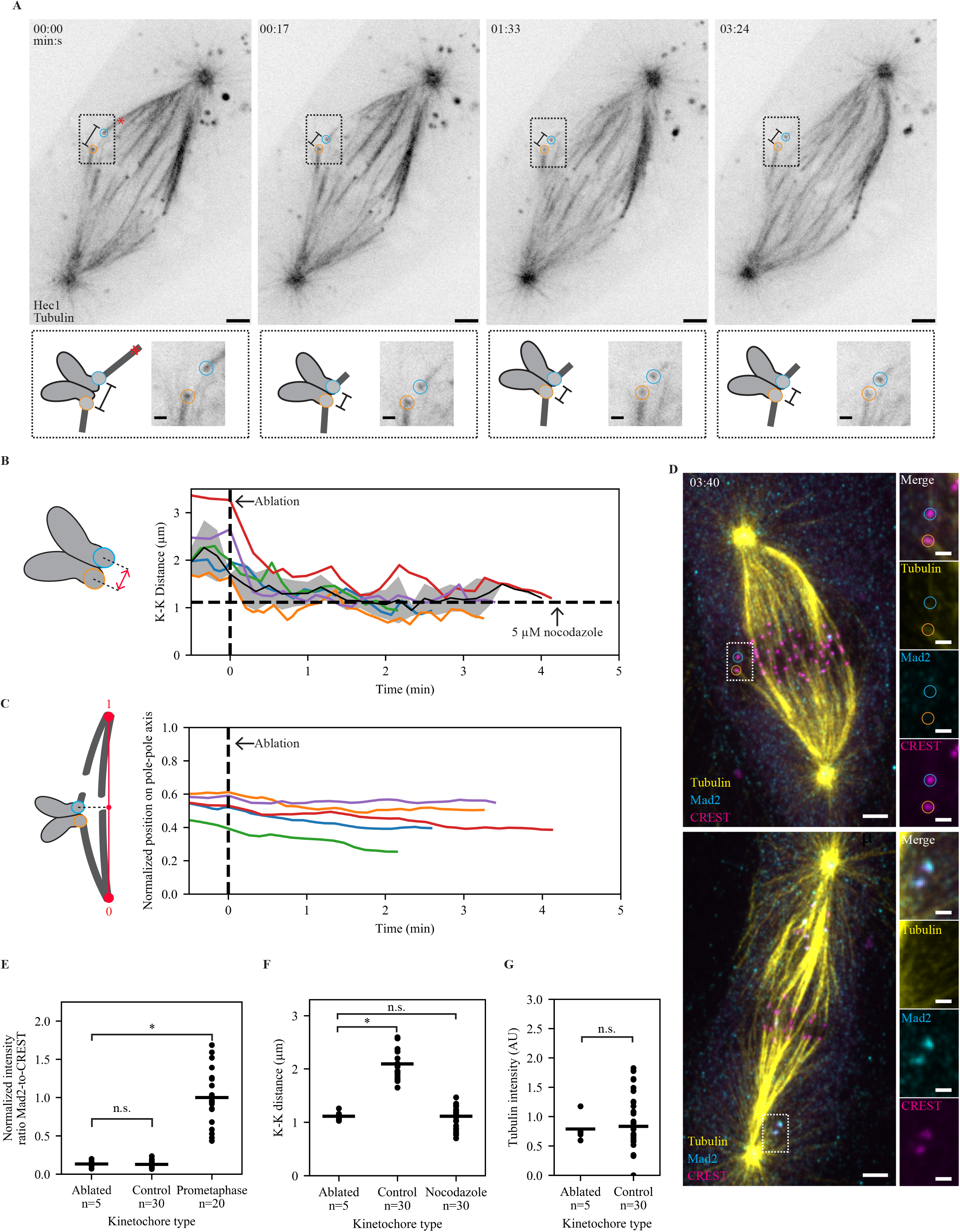
The mammalian SAC does not detect changes in spindle pulling forces. **(A)** Timelapse imaging (maximum intensity projection) of microtubule attachments (EGFP-tubulin) and kinetochores (Hec1-EGFP) in a metaphase Hec1 RNAi + partial NuMA RNAi PtK2 cell, during the mechanical isolation of the highlighted k-fiber (circles) using laser ablation (red X, t=0). Bottom: schematic and zoom of highlighted pair. Scale bars=3 μm (large) and 1 μm (zoom). **(B**) Mean, SEM, and individual K-K distance of pairs before and after ablation (time of fixation is approximately 30 s from the end of trace). Vertical dashed line marks first ablation. Horizontal dashed line marks the average K-K distance in 5 μM nocodazole (n=30 kinetochores). Example in (A) is the purple trace. **(C)** Normalized distance along the pole axis for disconnected kinetochores before and after ablation. Dashed line marks first ablation. Example in (A) is the purple trace. **(D)** Immunofluorescence imaging (maximum intensity projection) of microtubule attachment (tubulin), kinetochores (CREST), and SAC activation (Mad2) at (top) the cell in (A) and (bottom) a prometaphase cell on the same dish at approximately t=3:40. Scale bars=3 μm (Large) and 1 μm (zoom). **(D)** Individual (circles) and average (lines) SAC activation (Mad2/CREST, normalized to prometaphase intensity) at the ablated kinetochore (n=5), intracellular controls (n=30), and prometaphase cells on the same dish (n=20). There is no SAC activation on sister kinetochores attached to an ablated k-fiber. (**E)** Individual (circles) and average (lines) K-K distance at ablated, intracellular control, and nocodazole-treated pairs from (B). Ablation reduces K-K distance to a value similar to that in nocodazole (p=0.40). **(F)** Microtubule attachment intensity at the ablated and intracellular control pair. There is no difference in attachment number. (*; p<0.005, n.s; p>0.05, 2-sided Mann-Whitney U test).

To assess SAC signaling after prolonged loss of force, we fixed each ablated cell (3-5 min after the initial cut), stained for Mad2, kinetochores, and tubulin (Fig. 1D, top) and re-imaged each cell. There was no increase in kinetochore Mad2 intensity with an ablated k-fiber versus controls with no ablation in the same cell (Fig. 1D, top, and 1E, p=0.45), while unattached kinetochores in nearby cells had higher Mad2 intensity (Fig. 1D, bottom, and 1E, p=10^-4^). Ablating k-fibers led to a decrease in K-K distance compared to control pairs in the same cell (Fig. 1F, 1.12±0.04 vs 2.09±0.05 μm, p=10^-4^), but indistinguishable from pairs in nocodazole-treated cells (1.12±0.04 vs 1.11±0.04 μm, p=0.40). We also did not detect loss in k-fiber microtubule intensity after ablation (Fig. 1G, p=0.42), implying that the timescale of any force-based microtubule destabilization is longer than the time we allow. While these kinetochores do not re-recruit Mad2 despite losing spindle pulling forces (Fig. 1B-C, F), nocodazole-treated kinetochores re-recruit Mad1 on the same timescale as microtubule loss (Fig. S1). We conclude that the SAC does not detect changes in spindle pulling forces – even in the scenario of a k-fiber isolated from the spindle. Thus, microtubule binding itself controls SAC satisfaction at individual kinetochores.

After eliminating spindle forces as a confounding variable, we asked how microtubule attachment numbers are converted into a SAC response. Previous work suggested that Mad1 loss begins at about half of metaphase microtubule occupancy (Kuhn and Dumont, 2017). However, the decision to satisfy the SAC could have occurred before Mad 1 loss began. To test how steady-state microtubule attachment numbers affect SAC signaling, we reduced them by expressing a Hec1 variant (Hec1-9D-FusionRed) with greatly reduced microtubule affinity (Guimaraes et al., 2008; Zaytsev et al., 2015, 2014) in PtK2 Hec1 siRNA cells, and monitored EYFP-Mad1 loss dynamics during attachment formation (visualized using SiR-Tubulin (Lukinavičius et al., 2014)). While the Mad1 loss rate does not increase with higher attachment number (Kuhn and Dumont, 2017), we find that it decreases with decreased numbers (Fig. 2A-C, t_1/2_ = 190 s in Hec1-9D vs 79 s in wildtype, Video 2). A decrease in K-K distance versus wildtype (Fig. 2D, 1.49±0.03 vs 1.90±0.08 μm, p = 10^-6^) confirms a reduction in kinetochore-microtubule affinity and attachment number in Hec1-9D cells (Guimaraes et al., 2008). We conclude that the kinetochore responds to reduced microtubule levels by tuning the dynamics of its SAC response. While we still detected complete loss of Mad1 (Fig. 2B), the decreased rate of a Mad1 loss may impede satisfaction on weak attachments, allowing time for attachment reinforcement or error correction.

**Figure 2:**
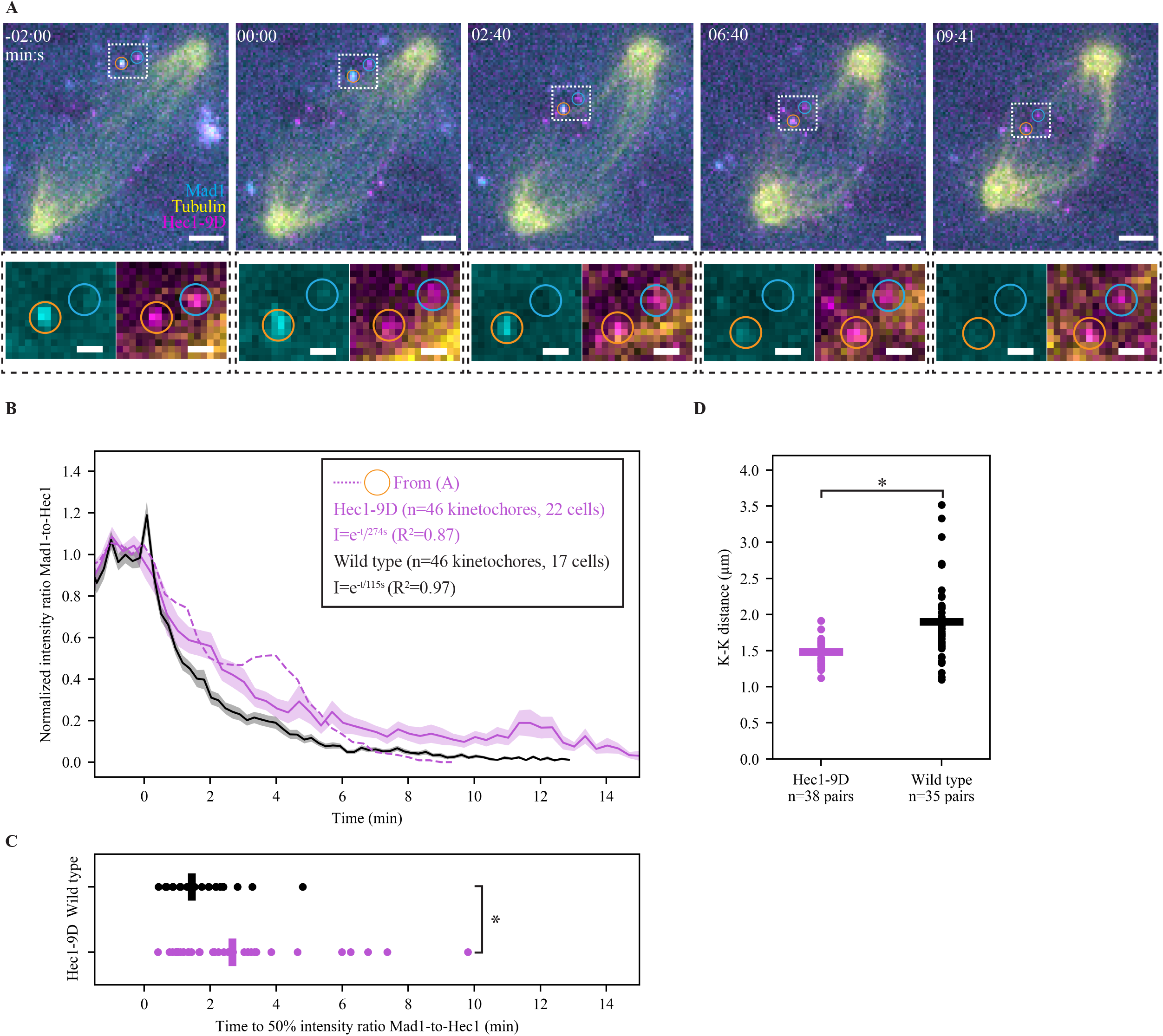
Lowering microtubule occupancy at a kinetochore slows down SAC satisfaction kinetics. **(A)** Timelapse imaging (maximum intensity projection) of representative SAC satisfaction kinetics (EYFP-Mad1) and microtubule attachment (SiR-Tubulin) in a Hec1-RNAi PtK2 cell with decreased spindle forces and kinetochore-microtubule occupancy (Hec1-9D-FusionRed). Scale bars=3 μm (large) and 1 μm (zoom), and t=0 indicates the start of Mad1 loss on the orange-circled kinetochore. **(B)** Mean, SEM, and individual trace of the orange-circled kinetochore in (A) of the Mad1-to-Hec1 intensity ratio with t=0 Mad1 loss start, and **(C)** distribution of times to reach a 0.5 intensity ratio of Mad1-to-Hec1, in wildtype (n=46 kinetochores) and in Hec1-9D-expressing (n=46) cells. Reducing steady-microtubule occupancy lowers Mad1 loss rates. Wildtype data taken from Kuhn and Dumont 2017, JCB. **(D)** Individual (circles) and average (lines) K-K distance in wildtype (n=35 pairs) and Hec1-9D (n=38) cells. Tension is reduced in Hec1-9D vs wild type cells. (*; p<0.005, 2-sided Mann-Whitney U test).

To quantify how microtubule attachment number and the SAC response scale with Hec1-microtubule binding levels, we developed a system to control the number of Hec1-microtubule interactions in an inducible Hec1 knockout HeLa cell line (Hec1Δ cells) (McKinley and Cheeseman, 2017). We reasoned that Hec1 RNAi + Hec1-9D cells (Fig. 2) may still satisfy the SAC and have a significant K-K distance due to residual Hec1-WT, and turned to Hec1 knockout cells. By deleting endogenous Hec1 (Fig. S2A-B) and co-expressing weak-binding Hec1-9D-FusionRed and metaphase-like affinity Hec1-1D8A-EGFP (Hec1-1D) (Zaytsev et al., 2014) (Fig. 3A), we created kinetochores with a variable mixture of Hec1 molecules with different affinities, indicated by the relative amounts of EGFP and FusionRed. Unlike the use of intermediate Hec1 affinity mutants, this approach limits the maximum number of Hec1 molecules bound to microtubules directly (Fig. 3A), without tuning the lifetime or affinity of individual Hec1-microtubule interactions, which could alter their ability to signal to the SAC (Hiruma et al., 2015; Ji et al., 2015). To probe the relationship between the functional Hec1 number and microtubule attachment number, we incubated Hec1Δ cells in proteasome inhibitor MG132 for 1 hr to allow kinetochores to reach steady-state attachment number and then fixed and stained for microtubules, EGFP, and FusionRed (Hec1-9D). As expected, high Hec1-1D cells formed robust microtubule attachments and high Hec1-9D cells did not (Fig. 3B, S2C). As the fraction of Hec1-1D increased, attachments produced more force as measured by the K-K distance (Fig. 3C, Spearman’s rho=0.35 p=10^-21^), and the intensity of end-on microtubule attachments rose in a graded, analog way (Fig. 3D, Spearman’s rho=0.38, p=10^-26^). Thus, this approach (the “mixed kinetochore” system) allows us to tune and fix kinetochore-microtubule numbers in a graded and quantifiable way, and reveals weak attachment cooperativity between Hec1 subunits in the *in vivo* native kinetochore, similar to *in vitro* measurements (Cheeseman et al., 2006; Ciferri et al., 2008; Zaytsev et al., 2015).

**Figure 3:**
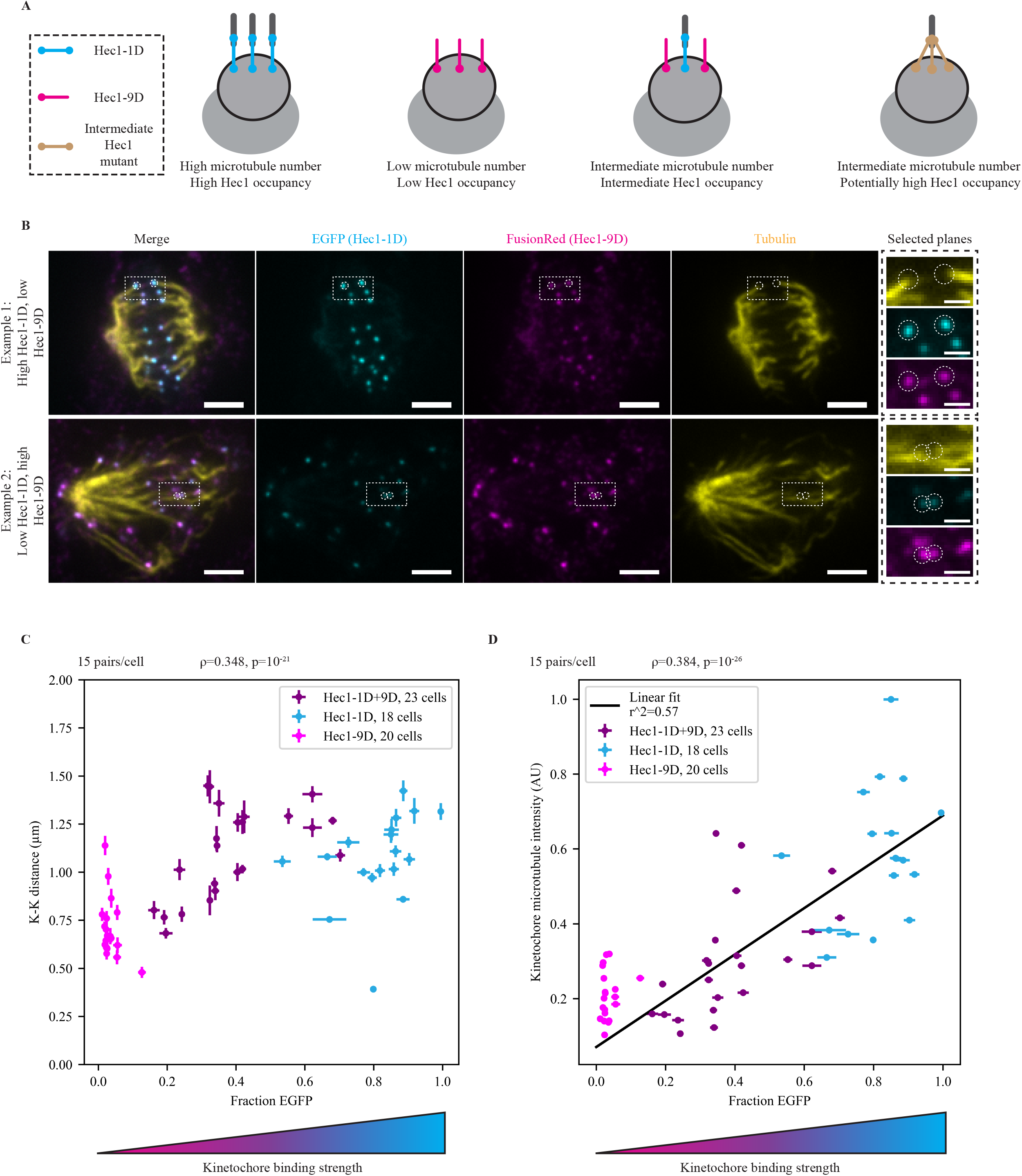
Kinetochore-microtubule occupancy scales with the number of functional Hec1 subunits in a smooth, analog fashion. **(A)** Schematic depicting experimental design. After deleting endogenous Hec1, strong (Hec1-1D, blue) and weak (Hec1-9D, pink) microtubule-binding mutants are expressed. Cells randomly receive different numbers of functional binders, and therefore have different microtubule occupancy. Unlike the expression of a single intermediate mutant, this approach limits the maximum number of Hec1 molecules on a kinetochore which can bind microtubules. **(B)** Immunofluorescence imaging (maximum intensity projection) of microtubule attachments (tubulin), Hec1-1D intensity (anti-EGFP), and Hec1-9D intensity (anti-mKate, binds to FusionRed) in Hec1Δ cells expressing Hec1-1D-EGFP and Hec1-9D-FusionRed. Cells were treated with 5 μM MG132 to accumulate cells at a metaphase spindle steady-state. The two highlighted examples were taken from the same coverslip, where the top has a high Hec1-1D to -9D ratio and the bottom a low ratio. Scale bars=3 μm (large) and 1 μm (zoom). **(C-D)** Mean of cellular EGFP fraction for each cells vs mean cellular K-K distance **(C)** and mean cellular kinetochore microtubule intensity **(D)** from cells in (B) (n=345 pairs, 690 kinetochores, 23 cells) and control 1D-(n=270, 540, 18) and 9D-alone coverslips (n=300, 600, 20) (Fig. S2C). Error bars=SEM.

We then used this mixed kinetochore system to map how kinetochores with different Hec1-microtubule binding levels coordinate a SAC output response (Fig. 4A). It is possible that every Hec1 molecule in the proposed outer kinetochore “lawn” (Zaytsev et al., 2014) functions independently, reducing Mad1 levels in a gradual, analog fashion as more Hec1s bind microtubules (Fig. 4A, top). Alternatively, the kinetochore could integrate information from all Hec1 binding sites and set a threshold fraction of Hec1s attached where the SAC turns off in a digital fashion (Fig. 4A bottom). To distinguish between these scenarios, we treated Hec1Δ cells expressing Hec1-9D or Hec-1D or varying mixtures of both with MG132 for 1 hr and then fixed and stained for Mad1, GFP (Hec1-1D), and FusionRed (Hec1-9D) (Fig. 4B). Cells with high Hec1-9D had many more Mad1-positive kinetochores than cells with high Hec1-1D (Fig. 4B-C, S3). Using the relationship between the Hec1-1D (EGFP) fraction and microtubule intensity in parallel experiments (Fig. 3D), we estimated that the lowest fraction EGFP mixed kinetochores (0.14) have ^~^22% of wildtype microtubule levels – about four attached microtubules (Wendell et al., 1993) (Fig. S3B). At these levels, many kinetochores had no detectable Mad1 (Fig. 4C, ^~^60%), suggesting that the increase in Mad1-positive kinetochores at low Hec1-9D reflects a change in kinetics (Fig. 2) rather than ability to satisfy the SAC. We conclude that approximately four microtubules are sufficient for SAC satisfaction. SAC satisfaction with few microtubule attachments and many Hec1s that cannot bind microtubules subunits implies that the kinetochore integrates attachment information and does so in a switch-like manner (Fig. 4A bottom). Taken together, our data indicate that the mammalian kinetochore exclusively monitors microtubule attachment, and responds as a single unit even with few kinetochore-microtubules and a reduced number of bound Hec1s (Fig. 5).

**Figure 4:**
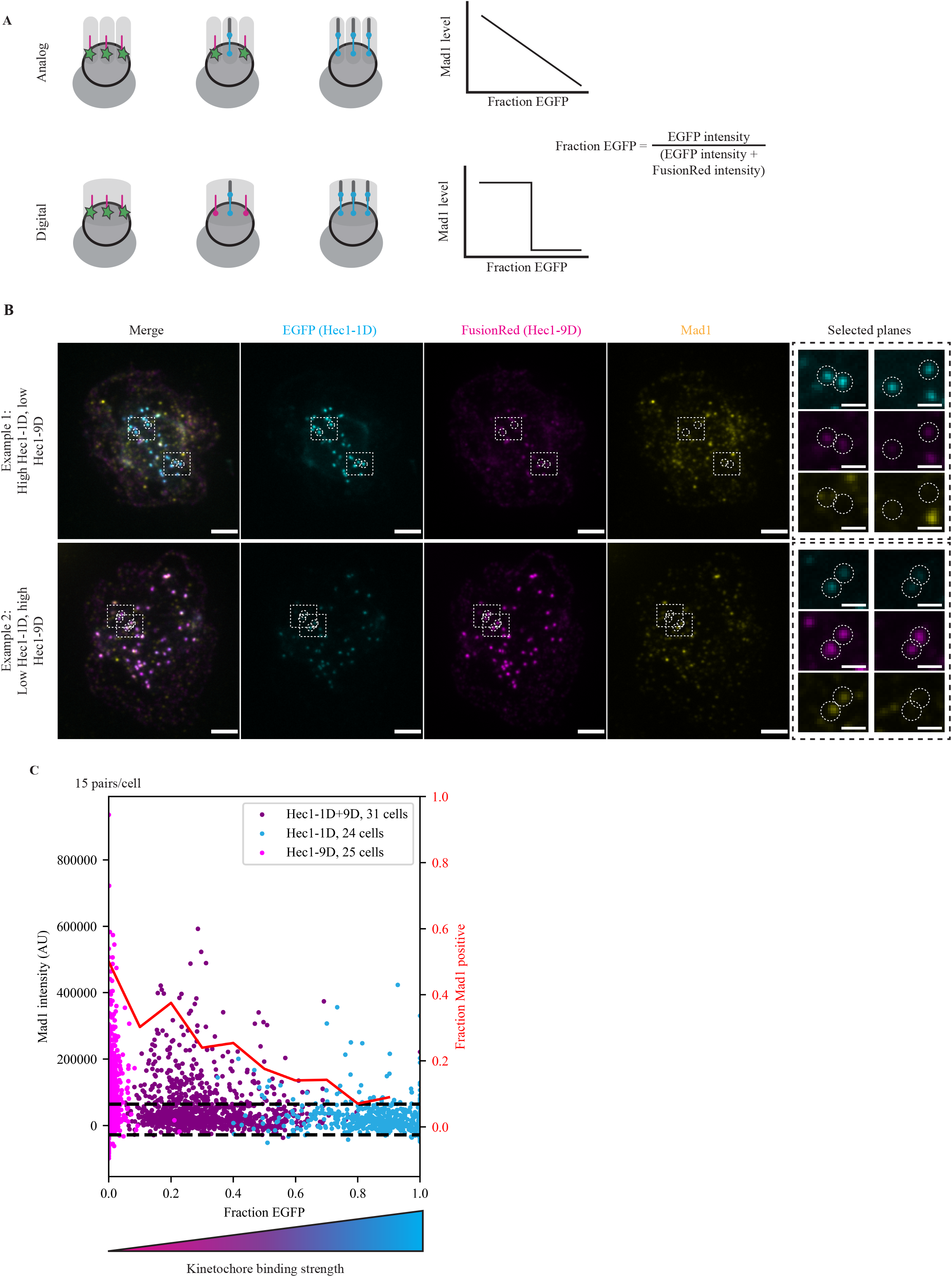
The kinetochore responds to the number of attached microtubules in a switch-like, highly sensitive manner. **(A)** Schematic depicting models for kinetochore signal integration. The kinetochore either processes microtubule attachments as many individual units (light grey boxes) and responds to attachments in an analog manner (top) or as one single unit and responds digitally (bottom). **(B)** Immunofluorescence imaging (maximum intensity projection) of SAC activation (Mad1), Hec1-1D intensity (anti-EGFP), and Hec1-9D intensity (anti-mKate, binds to FusionRed) in Hec1Δ cells expressing both Hec1-1D-EGFP and Hec1-9D-FusionRed. Cells were treated with 5 μM MG132 to accumulate cells at a metaphase spindle steady-state. The two examples are cells on the same coverslip where the top has a high Hec1-1D to -9D ratio and the bottom a low ratio. Kinetochores in both conditions are capable of recruiting (left zoom) and losing (right zoom) Mad1. Scale bars=3 μm (large) and 1 μm (zoom). **(C)** Fraction EGFP vs Mad1 intensity from cells in (B) (n=930 kinetochores, 31 cells) and 1D-(n=720, 24) and 9D-(n=750, 25) alone control cells (Fig. S3A). Red line indicates the fraction of kinetochores with Mad1 intensities one standard deviation (horizontal lines) greater than average Mad1 intensity on Hec1-1D kinetochores.

**Figure 5:**
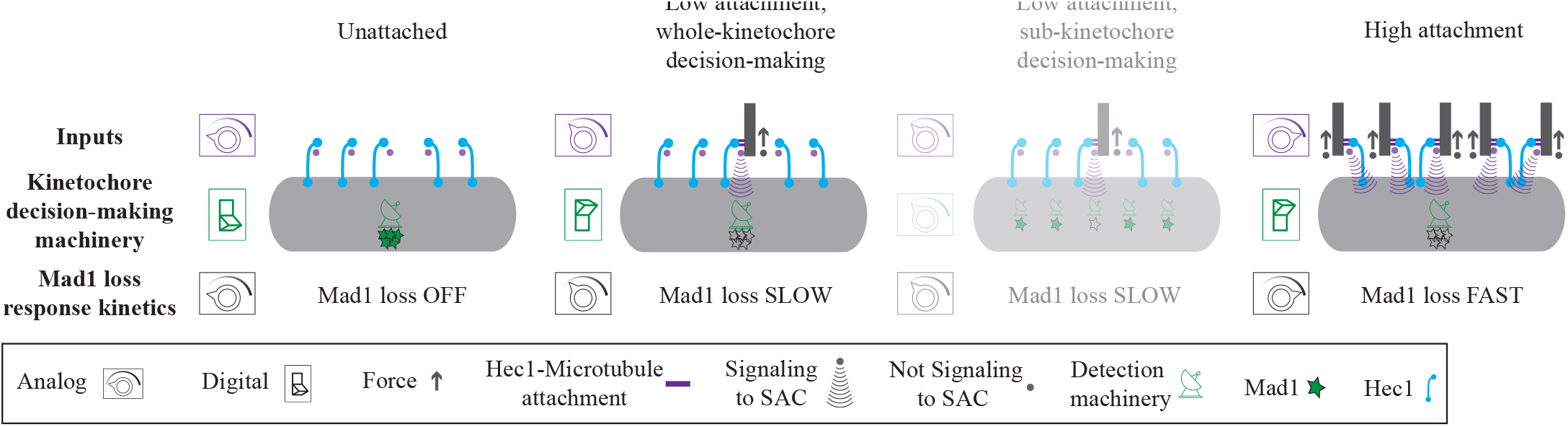
The mammalian kinetochore integrates attachment signals in a sensitive and switch-like manner. Potential SAC inputs (Hec1-microtubule binding and spindle forces) are analog, varying widely in time and between kinetochores (top). We show that the SAC decision-making machinery (middle) responds specifically to Hec1-microtubule binding and activates Mad1 removal machinery (bottom). We further show that in contrast to its analog inputs, SAC decision-making is digital: even kinetochores with very few microtubule attachments initiate a full Mad1 loss response, indicating a form of whole-kinetochore signal amplification and integration, and inconsistent with sub-kinetochore decision-making (shaded cartoon). Downstream of decision-making, the kinetochore is capable of tuning Mad1 loss rates in response to reduced attachment. The combination of digital decision-making and an analog response rate is well-suited to allow cells to rapidly exit mitosis while preventing errors (see Discussion).

## Discussion

In the SAC, the kinetochore acts as a subcellular computer: it accepts inputs from the environment, processes them, and produces an output controlling cell cycle progression. While we are defining the necessary attachment inputs, the outputs (Mad1 localization), and the kinetochore “hardware” (kinetochore structure and SAC biochemistry), we know little about its “software”: how different inputs are sensed and how inputs and outputs are quantitatively related. Here, we show that the kinetochore is not sensitive to mechanical force (Fig. 1). Instead, the kinetochore is programmed to respond to microtubule attachment number in two ways (Fig. 5): the magnitude of the SAC response (rate of Mad1 loss) scales with the level of microtubule attachment (Fig. 2), and the kinetochore decision-making process (initiation of Mad1 loss) is digital, where the SAC is satisfied with four microtubules (Fig. 3-4).

The link between mechanical force and kinetochore-microtubule attachment inputs has made dissecting how they modulate SAC activity challenging. K-fiber laser ablation (Fig. 1) allowed us to remove spindle pulling forces without perturbing global microtubule dynamics or kinetochore proteins. As such, our findings provide the most direct demonstration that the SAC does not detect spindle-generated forces at individual kinetochores. If the intra-kinetochore deformations which may be involved in SAC signaling (Maresca and Salmon, 2009; Uchida et al., 2009) persist after ablation, our data shows they do not reflect force levels relevant for kinetochore mechanical function. Recently, it was observed that tensionless, attached kinetochores activate the SAC after depletion of Kif18A or Astrin (Janssen et al., 2018). It is possible that either depleting these proteins alters kinetochore-microtubule attachments (Shrestha et al., 2017; Stumpff et al., 2012) or SAC function, or that spindle forces are needed for establishing but not maintaining SAC satisfaction. Notably, unlike force, plus-end microtubule attachments only stably exist in bioriented kinetochores (King and Nicklas, 2000). Therefore, for kinetochores to monitor microtubule occupancy may enhance the SAC’s reliability.

The removal of Mad1 from individual kinetochores is likely controlled by one rate-limiting step (Kuhn and Dumont, 2017). By reducing the number of kinetochore-bound microtubules, we show that this process (downstream of decision-making) is tuned by microtubule number (Fig. 2). This may be related to microtubules’ role as tracks for dynein transport of Mad1/2 from the kinetochore (Howell et al., 2001): reducing the number of dynein tracks could reduce SAC protein stripping kinetics. Previously, neither changing centromere tension nor increasing attachment number affected Mad1 loss rates (Kuhn and Dumont, 2017), suggesting that wildtype kinetics are limited by dynein concentration or kinetochore biochemistry. Linking Mad1 loss rates to attachment status may prevent rapid satisfaction on incorrect attachments. This is critical because microtubule detachment (error correction) at low force is slow (Fig. 1) and the SAC begins to satisfy with few kinetochore-bound microtubules (Fig 4).

Using mixed kinetochores, we probed how the disordered kinetochore “lawn” works as an ensemble to control k-fiber structure and kinetochore decision-making. We found that the number of k-fiber microtubules scales gradually with functional Hec1 (Fig. 3), indicating a lack of cooperativity which may prevent reinforcement of incorrect attachments. In contrast, the relationship between microtubule binding and the SAC is switch-like and sensitive (Fig. 4): many satisfied kinetochores in our assay likely cannot reach metaphase levels of microtubule-bound Hec1 (32%) (Yoo et al., 2018). The response to low attachment number requires integrating attachment information over the kinetochore (Fig. 5). Feedback between SAC kinases and phosphatases could amplify small decreases in kinase activity, creating a switch-like response (Funabiki and Wynne, 2013; Nijenhuis et al., 2014; Saurin et al., 2011). There are two recent models for how microtubule binding could reduce kinase activity: occupying Mps1 binding sites (Hiruma et al., 2015; Ji et al., 2015) or displacing kinases from substrates (Aravamudhan et al., 2015; Hengeveld et al., 2017). In a competition model, the signal amplification we observe explains how metaphase kinetochores with highly variable attachment number and some Mps1 localization (Howell et al., 2004) lose Mad1; in a displacement model, our data suggest that displacement does not reflect relevant changes in mechanical force.

The low number of kinetochore-microtubules required for SAC satisfaction not only has important implications for how the kinetochore “computes”, but also for how it functions in a cellular context. For example, this low number may prevent high variability in kinetochore-microtubule numbers (McEwen et al., 1997) from causing mitotic delays which lead to DNA damage and cell death (Orth et al., 2012; Uetake and Sluder, 2010). Looking forward, probing how the number of required kinetochore-microtubules changes after modifying kinetochore structure and biochemistry will be critical to understanding how the kinetochore processes physical information into a robust signaling biochemical response.

## Materials and Methods

### Cell culture and transfection

PtK2 EYFP-Mad1 (Shah et al., 2004) (gift from Jagesh Shah) and wild type cells were cultured in MEM (Thermo Fisher, Waltham, MA) supplemented with sodium pyruvate (11360; Thermo Fisher), nonessential amino acids (11140; Thermo Fisher), penicillin/streptomycin, and 10% heat-inactivated fetal bovine serum (FBS) (10438; Thermo Fisher). Tet-on inducible CRISPR-Cas9 HeLa cells (Hec1Δ cells, gift from Iain Cheeseman, Whitehead Institute) were cultured in DMEM/F12 with GlutaMAX (10565018; Thermo Fisher) supplemented with penicillin/streptomycin, 5 μg/mL puromycin, and tetracycline-screened FBS (SH30070.03T; Hyclone Labs, Logan, UT). Cas9 expression was induced by the addition of 1 μM doxycycline hyclate 48 hr before fixation. Knockout was confirmed by Western Blot and the accumulation of mitotic cells after doxycycline addition. Cell lines were not STR-profiled for authentication. All cell lines tested negative for mycoplasma. Cells were maintained at 37°C and 5% CO_2_. For imaging, cells were plated on 35 mm #1.5 glass-bottom dishes (poly-d-lysine coated, MatTek, Ashland, MA; Fig. 2, S1), 25 mm # 1.5 glass etched coverslips (acid cleaned and poly-L-lysine coated; G490; ProSciTech, Kirwan, AUS; Fig. 1), or 25 mm # 1.5 glass coverslips (acid cleaned and poly-L-lysine coated; 0117650; Marienfeld, Lauda-Königshofen, DEU; Fig. 3, 4 S2, S3). Cells were transfected with EGFP-Tubulin (Clontech, Mountain View, CA, discontinued), Hec1-WT-EGFP (gift from Jennifer DeLuca, Colorado State University), mCherry-CenpC (gift from Aaron Straight, Stanford University), Hec1-8AS8D-EGFP (1D-EGFP; gift from J. DeLuca, Colorado State University), and Hec1-9D-FusionRed (FusionRed (gift from Michael Davidson) was swapped for EGFP in Hec1-9D-EGFP (gift from Jennifer DeLuca, Colorado State University)) using ViaFect (E4981; Promega, Madison, WI) 48 hr (HeLa) or 72 hr (PtK2) before imaging. For siRNA knockdown, cells were treated with 100 nM siRNA oligos (Sigma-Aldrich, St. Louis, MO) and 4 μl oligofectamine (12252011; Thermo Fisher) either 6 hr (Fig. 1) or 24 hr (Fig. 2) after transfection (66 or 48 hr before imaging, respectively). The following oligos were used: siNuMA: 5′-GCATAAAGCGGAGACUAAA-3′ (Elting et al., 2017); siHec1 5’-AATGAGCCGAATCGTCTAATA-3’ (Guimaraes et al., 2008).

### Immunofluorescence

For “ablate and fix” experiments (Fig. 1), cells were fixed in 95% methanol + 5 mM EGTA for 1 min on ice. Cells were then blocked at room temperature for 1.5 hr in TBST (50 mM Tris, 150 mM NaCl, 0.05% Triton X100, pH 7.6) + 2% BSA. Primary and secondary antibody incubations were done in blocking solution for 1 hr and 30 min, respectively. Between each step, 4-5 washes in TBST were performed. For “mixed kinetochore” immunofluorescence (Fig. 3, 4, S2, S3), cells were pre-extracted in PHEM (120 mM PIPES, 50 mM HEPES, 20 mM EGTA, 4 mM Magnesium Acetate, pH 7) + 1% Triton X-100 for 20 s then fixed for 15 min in PHEM + 4% PFA (freshly dissolved from powder) at 37°C. Cells were then permeabilized in PHEM + 0.5% IGEPAL-CA-630 for 10 min and blocked in PHEM + 0.05% Triton X-100 + 2% BSA for 1 hr at room temperature. Both the primary and secondary antibody incubations were done in block solution at 37° for 1 hr. Between each step (excluding pre-extraction), 4-5 min washes in PHEM + 0.05% Triton X-100 were performed; PFA staining protocol adapted from (Suzuki et al., 2018). For all experiments, cells were mounted in ProLong Gold (P10144; Thermo Fisher) and stored in the dark at 4°C. The following antibodies were used: mouse anti-α-tubulin DM1 (1:1,000; T6199; Sigma-Aldrich), human anti-centromere (CREST; 1:25; discontinued; Antibodies Inc., Davis, CA), rabbit anti-rat kangaroo-Mad2 (DeLuca et al., 2018; Kuhn and Dumont, 2017) (1:100), mouse anti-Hec1 (9G3; 1:100, ab3613; Abcam), rabbit anti-mKate (recognizes FusionRed) (1:200; TA150072; OriGene, Rockville, MD), mouse anti-hsMad1 (1:300; MABE867; EMD-Millipore, Burlington, MA), rabbit anti-alpha-tubulin (1:200; ab18251; Abcam, Cambridge, UK), camel anti-EGFP conjugated to Atto488 (1:100; added during secondary; gba-488; ChromoTek, Hauppauge, NY), anti-mouse secondary antibodies (1:500) conjugated to Alexa-488 (A11001; Invitrogen, Carlsbad, CA) or Alexa-647 (A21236; Invitrogen), anti-rabbit secondary antibodies (1:500) conjugated to Alexa-488 (A11008; Invitrogen), Alexa-568 (A11011; Invitrogen) or Alexa-647 (A21244; Invitrogen), and a human secondary antibody conjugated to DyLight 405 (1:100; 109-475-098; Jackson ImmunoResearch Laboratories, West Grove, PA).

### Drug and dye treatments

To depolymerize spindle microtubules (Fig. 1, S1), 5 μM nocodazole (M1404; Sigma-Aldrich) was added 10 min prior to fixation (Fig. 1) or 10 μM was added at the indicated time (Fig. S1). To prevent anaphase onset (Fig. 3,4, S2, S3) cells were treated with 5 μM MG132 (474790; EMD-Millipore) 1 hr prior to fixation. To visualize tubulin as a third color (Fig. 2), 100 nM SiR-Tubulin dye (cy-sc002; Cytoskeleton, Inc., Denver, CO) was added 1 hr prior to imaging, along with 10 μM verapamil (V4629; Sigma-Aldrich) to prevent dye efflux.

### Imaging

#### Microscope settings

All imaging was performed on an inverted (Eclipse Ti-E; Nikon, Tokyo, JPN), spinning disk confocal (CSU-X1; Yokogawa Electric Corporation, Tokyo, JPN) microscope. Single color live imaging (Fig. 1) was performed with a Di01-T488-13×15×0.5 (Semrock, Lake Forest, IL) head dichroic along with a 488 nm (120 mW) diode laser, an ET500LP emission filter (Chroma, Bellows Falls, VT), and an iXon3 camera (Andor Technology, Belfast, UK; bin=1, 105 nm/pixel). For these experiments, cells were imaged in phase contrast (400 ms exposure) and fluorescence (60 ms exposure) in 3 z-planes spaced 700 nm apart every 7.5-15 s with a 100×1.45 Ph3 oil objective through a 1.5× lens with 5x pre-amplifier gain and no EM gain (Metamorph 7.7.8.0; Molecular Devices, San Jose, CA).

Three-color live imaging (Fig. 2, S1) was performed with a Di01-T405/488/568/647 head dichroic (Semrock) instead, along with 561 nm (150 mW) and 642 nm (100 mW) diode lasers and different emission filters (ET525/50M, ET630/75M, and ET690/50M; Chroma). Cells were imaged by phase contrast (200 ms exposure) and fluorescence (40-75 ms exposure) in four z-planes spaced 350 nm apart every 13-30 s and at bin=2 (to improve imaging contrast for dim Mad1 and microtubule structures; 210 nm/pixel). All live PtK2 cells were imaged at 30 °C, 5% CO2 in a closed, humidity-controlled Tokai Hit (Fujinomiya, JPN) PLAM chamber.

For fixed cell imaging (Fig. 1, 3, 4, S2, S3) a 405 nm (100 mW) laser was added along with an ET455/50M emission filter (Chroma) and two emission filters were changed to ET525/36M and ET600/50M (Chroma). Cell images were acquired in z-slices 300 nm apart with bin=1 and laser powers, exposure times, and EM Gain optimized (but not changed between cells) to fill as much of the dynamic range of the camera as possible without saturation. For mixed kinetochore experiments (Fig. 3, 4), all acquisition settings for EGFP and FusionRed were kept identical.

#### Ablation Protocol

Laser ablation (20 3-ns pulses at 20Hz) with 551nm light was performed using the MicroPoint Laser System (Photonic Instruments, Belfast, UK). Images were acquired more slowly prior to ablation and then acquired more rapidly after ablation (typically 7.5 s prior and 15 s after). Successful k-fiber ablation was verified by loss of tension across the centromere (Fig. 1). To prevent k-fiber reincorporation into the spindle (Elting et al., 2014; Sikirzhytski et al., 2014), the spindle area around the k-fiber was also ablated concurrently and the minus-end of the k-fiber was re-ablated periodically prior to fixation.

#### Cell selection

For laser ablation (Fig. 1), metaphase cells with minor pole-focusing defects and wavy spindle morphology, indicative of partial NuMA knockdown, and visible Hec1-EGFP expression were chosen. In addition, Hec1 knockdown was confirmed by the lack of k-fibers and irregular motion of chromosomes in EGFP-negative cells. For imaging Mad1 loss (Fig. 2), prometaphase cells with moderate Mad1-EFYP expression, high Hec1-9D-FusionRed expression, and low average K-K distances (to indicate lack of strong attachments) were chosen. Hec1 knockdown was confirmed by the lack of k-fibers and irregular motion of chromosomes in FusionRed-negative cells.

### Data Analysis

#### Tracking and feature identification

For live ablation experiments (Fig. 1), kinetochores (Hec1-EGFP) and poles (EGFP-tubulin) were tracked by hand using a custom-made Matlab (Mathworks, Natick, MA) GUI (available here: https://github.com/DumontLab/Image-Analysis-Gui). Pairs were then included in further analysis if they exhibited prolonged decrease in K-K distance after ablation. For live imaging of Mad1 intensity (Fig. 2), kinetochores were tracked as previously (Kuhn and Dumont, 2017), using Matlab program SpeckleTracker (Wan et al., 2012). For analysis of fixed images (Fig. 1, 3, 4, S2, S3), kinetochores were identified by hand in a custom Matlab GUI using the plane of brightest Hec1 or CREST intensity and K-fibers were identified as bundles of tubulin intensity (where applicable).

#### Intensity measurements

Fixed kinetochore intensities (Fig. 1, 3, 4, S2, S3) were measured in Matlab by summing pixel intensities in a 7×7 (0.73×0.73 μm) box centered at the indicated coordinate. To calculate the Mad2/CREST ratio (Fig. 1) and Mad1 kinetochore intensity (Fig. 4), intensities were background-corrected by dividing (Fig. 1) or subtracting (Fig. 4) the kinetochore intensity by the average of three background intensities. To calculate the fraction EGFP (Fig. 3,4), the kinetochore EGFP intensity (background subtracted) was divided by the sum of the kinetochore EGFP and FusionRed intensity (both background subtracted). To calculate tubulin intensity on a given kinetochore, two 0.5 μm long intensity linescans were taken for each kinetochore: one (Tub_in_) perpendicular to the kinetochore-kinetochore axis 0.25 μm away from the kinetochore towards its sister, and one (Tub_out_) perpendicular to the kinetochore-microtubule axis 0.25 μm away from the kinetochore towards the microtubule. The microtubule attachment intensity is the difference between Tub_out_ and Tub_in_. To account for variance in staining between coverslips, all tubulin intensities were normalized to the intensity of a 7×7 pixel box centered on the spindle pole.

To determine Mad1 loss rates (Fig. 2), we measured EYFP-Mad1 and FusionRed-Hec1-9D intensities at each timepoint following a protocol identical to the one used to measure Mad1 loss rates previously (Kuhn and Dumont, 2017). In short, movies were thresholded by setting to zero all pixels <2 standard deviations above image background at the first frame. For each time point, the intensities of all pixels in a 5×5 pixel (1.05×1.05 μm) box around the kinetochore were summed together over all planes. t=0 was set to the time for each kinetochore where Mad1 intensity started decreasing while Hec1 intensity stayed constant, and intensities were normalized to the average intensity for t=-100 to t=0 (Kuhn and Dumont, 2017).

#### Statistics

Data are expressed as mean ± SEM. Calculations of p-Values (Mann-Whitney U) and correlation coefficients (Spearman rank-order) were done using Scipy and Numpy Python modules. To calculate the relationship between the fraction EGFP and the number of attached microtubules, a linear regression (least-squares) was applied to the data using Scipy. The lower limit of fraction of a metaphase attachment is calculated as the ratio between attachment numbers at the minimum fraction EGFP (average, mixed kinetochores) and the attachment number at fraction EGFP=1.

## Acknowledgements

We thank Jagesh Shah for EYFP-Mad1 PtK2 cells, Iain Cheeseman for Hec1Δ HeLa cells, Michael Davidson for the FusionRed construct, Aaron Straight for the mCherry-CenpC construct, Jennifer DeLuca for the Hec1-9D-EGFP, Hec1-WT-EGFP, and Hec1-1D-EGFP constructs, Jennifer DeLuca and Jeanne Mick for the rat kangaroo Mad2 antibody, Xiaohu Wan for Matlab SpeckleTracker, Eline ter Steege for preliminary experiments, David Morgan and Fred Chang for critical reading of the manuscript, and the Dumont lab for discussions and critical reading of the manuscript. We thank Geert Kops for sharing data prior to publication. This work was funded by NIH DP2GM119177 (S.D.), the Rita Allen Foundation and Searle Scholars’ Program (S.D.), the NSF Center for Cellular Construction 1548297 (S.D.), and a NSF GRF (J.K.). The authors declare no competing financial interests.

## Author contributions

J.K. developed methodology, performed experiments, analyzed the data and wrote the manuscript. J.K. and S.D. conceived the project and designed experiments. S.D. edited the manuscript.

---

**Figure S1:**
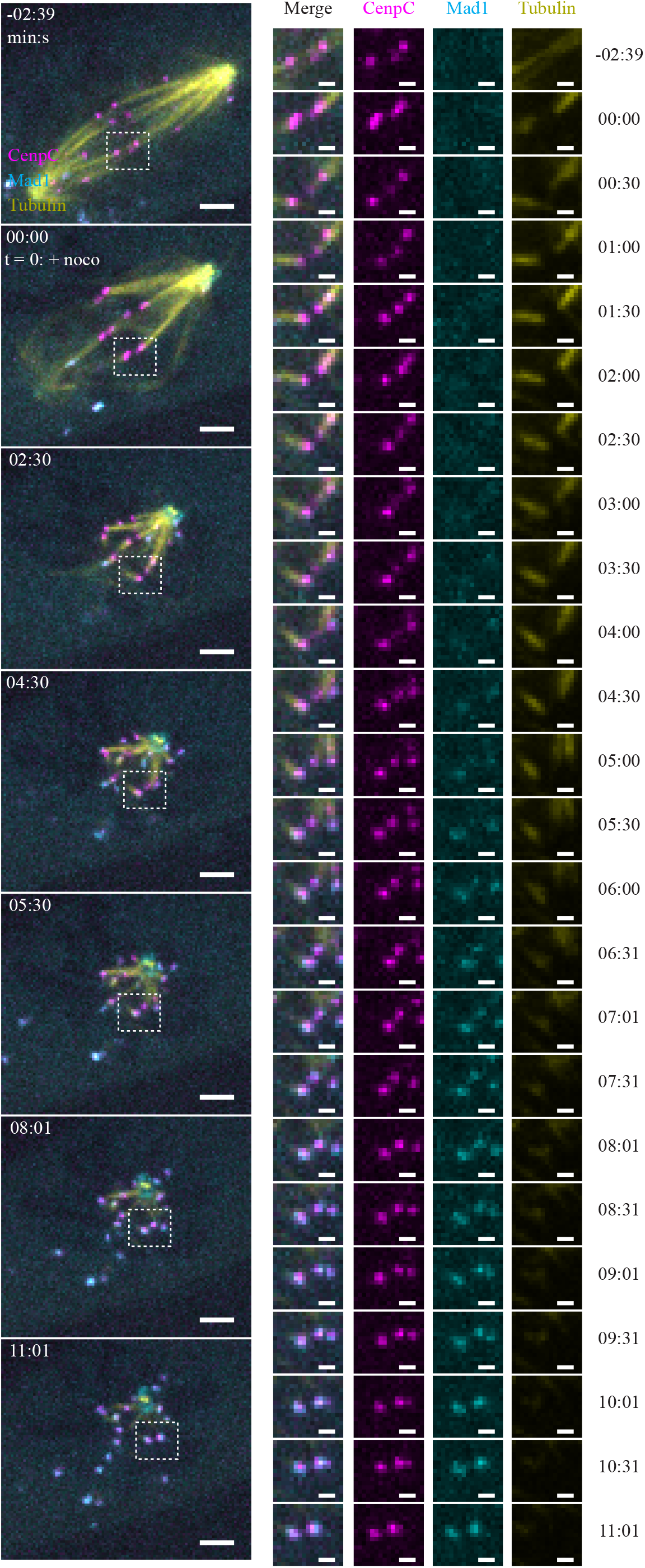
Mad1 is rapidly re-recruited upon nocodazole addition; related to Fig. 1. Timelapse imaging (maximum intensity projection) of representative SAC activation kinetics (EYFP-Mad1) and microtubule attachment (SiR-Tubulin) on kinetochores (CenpC-mCherry) in PtK2 cells after treatment with 10 μM nocodazole. Nocodazole addition (t=0) does not immediately depolymerize microtubule attachments. Mad1 is re-recruited on individual kinetochores rapidly only once attachments start to disappear. Scale bars=3 μm (large) and 1 μm (zoom).

**Figure S2:**
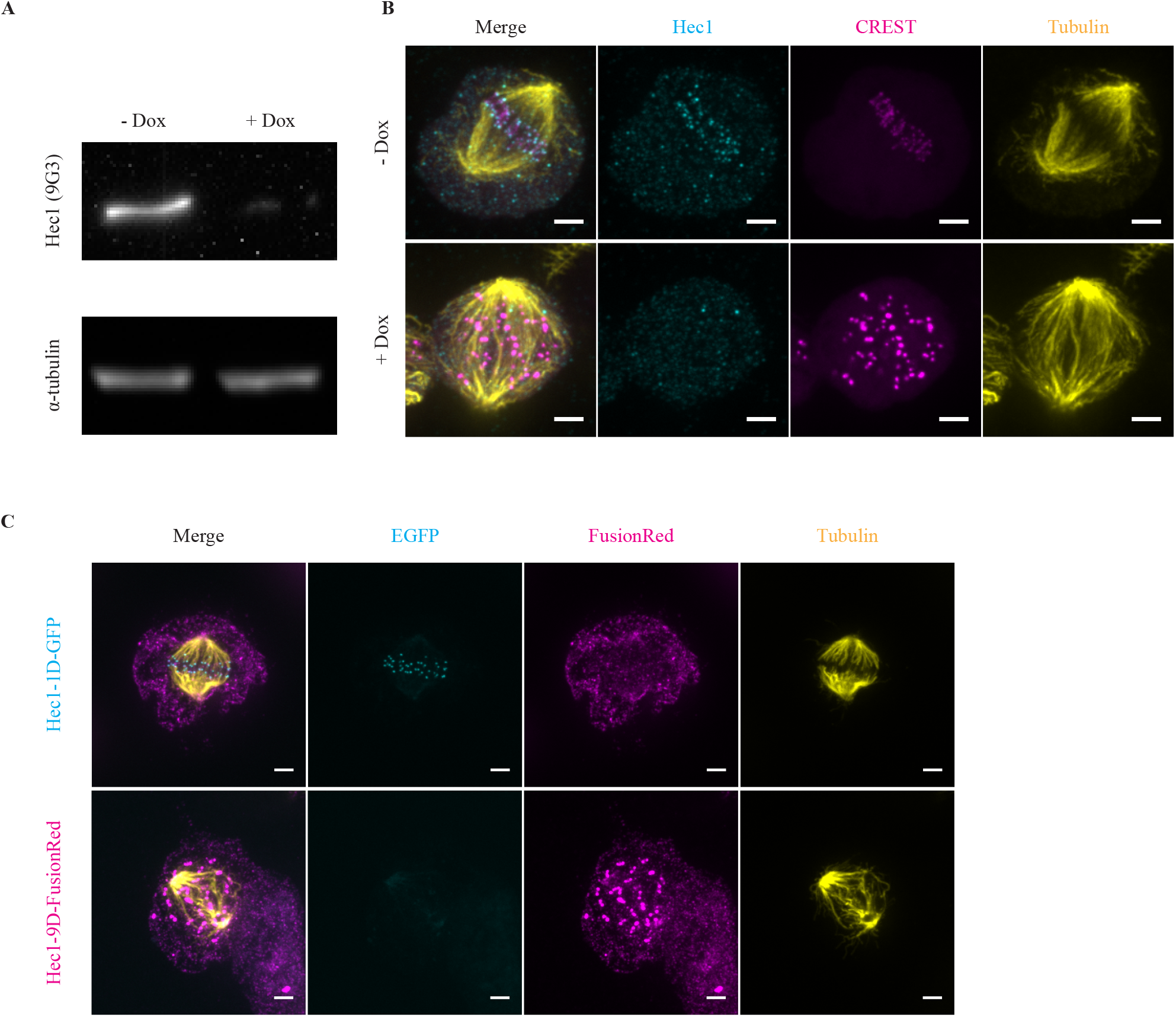
Hec1-1D, but not 9D, rescues spindle defects after Hec1 depletion; related to Fig. 3. **(A)** Western blot of Hec1 and tubulin abundance in cells with a stably-expressed doxycycline-inducible Cas9 and Hec1 sgRNA (Hec1Δ cells) (McKinley and Cheeseman, 2017). **(B)** Immunofluorescence imaging (maximum intensity projection) of microtubule attachments (tubulin), kinetochores (CREST) and Hec1 intensity in + and −dox Hec1Δ cells. Scale bars=3 μm. **(C)** Immunofluorescence imaging (maximum intensity projection) of microtubule attachments (tubulin), Hec1-1D intensity (anti-EGFP), and Hec1-9D intensity (anti-mKate, binds to FusionRed) in Hec1Δ +dox cells expressing Hec1-1D-EGFP of Hec1-9D-FusionRed. Cells were treated with 5 μM MG132 to accumulate cells at a metaphase spindle steady-state. Hec1-1D expression, but not-9D, rescues the spindle structure defects in (B).

**Figure S3:**
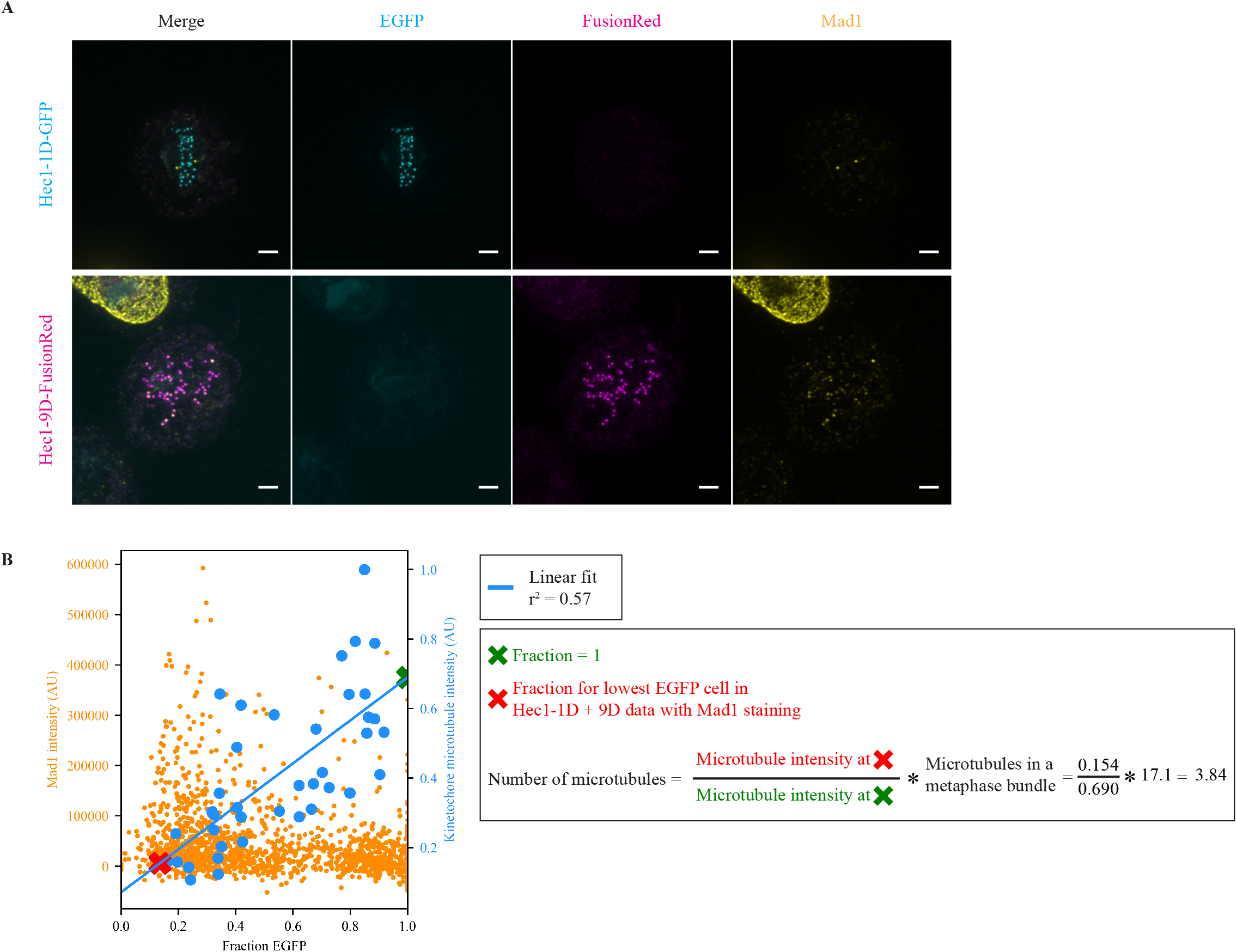
Hec1-1D, but not 9D, allows for consistent SAC satisfaction after Hec1 depletion; related to Fig. 4. **(A)** Immunofluorescence imaging (maximum intensity projection) of SAC activation (Mad1), Hec1-1D intensity (anti-EGFP), and Hec1-9D-FusionRed intensity (anti-mKate, binds to FusionRed) in Hec1Δ +dox cells expressing Hec1-1D-EGFP or Hec1-9D-FusionRed. Cells were treated with 5 μM MG132 to accumulate cells at a metaphase spindle steady-state. Hec1-1D expression, but not -9D, allows SAC satisfaction on most kinetochores. Scale bars=3 μm. **(B)** Fraction EGFP vs Mad1 intensity (Fig. 4) or average end-on attachment number (Fig. 3B-D) for Hec1-1D, and Hec1-1D + Hec1-9D conditions. Blue line indicates linear fit for fraction EGFP vs kinetochore-microtubule intensity (r^2^=0.57, p=10^-8^). X’s indicate the points along the fit used for the calculation of attached microtubule number (red = lowest average cellular fraction EGFP in the Mad1 vs. fraction data, green = 100% EGFP). Calculation of the attachment numbers uses average number of microtubules attached to a HeLa cell at metaphase from Wendell et al., 1993.

**Video 1: Mechanical isolation of a kinetochore-fiber from a metaphase spindle.** Timelapse spinning disk confocal imaging (maximum intensity projection) of representative mechanical isolation of a kinetochore by severing its k-fiber (Hec1-EGFP and EGFP-Tubulin, black) at an aligned kinetochore pair in a PtK2 cell with NuMa partially depleted and endogenous Hec1 depleted by RNAi. Upon initial laser ablation (red asterisks, t=0), the kinetochore whose k-fiber was severed (cyan circle) recoils towards its sister (orange circle). After ablation, the K-K distance remained low and the ablated k-fiber remained isolated from other spindle microtubules, never moving back towards it former pole. Periodically, the free minus was re-ablated (red asterisks) to prevent reincorporation of the k-fiber into the spindle. The cell was fixed 16 s after the last frame. Three planes spaced 700 nm apart were acquired every 15 s (before ablation and periodically after) or 7.5 s (after ablation). On timepoints with 15 s intervals, duplicate frames were added. Playback is 10 frames/s. Scale bar=3 μm. Time is in min:s. Video corresponds to images from Fig. 1A,D.

**Video 2: Lowering microtubule occupancy at a kinetochore slows down SAC satisfaction kinetics.** Timelapse spinning disk confocal imaging (maximum intensity projection) of a representative kinetochore pair’s SAC satisfaction kinetics (EYFP-Mad1, green) and attached microtubules’ geometry (SiR-Tubulin, yellow) at kinetochores with reduced microtubule affinity (Hec1-9D-FusionRed, magenta) during spindle assembly in a PtK2 cell with endogenous Hec1 depleted by RNAi. Prior to t=0, one kinetochore is Mad1-negative and has a weak microtubule end-on attachment (cyan circle) while its sister is Mad1-positive and unattached (orange circle). Mad1 loss (orange arrow) on the orange-circled kinetochore begins at t=0 as the pair moves towards the metaphase plate, consistent with acquiring an end-on attachment. Mad1 remains detectable until t=7:26, significantly longer than in wild-type cells. Four planes spaced 350 nm apart were imaged every 20 s. Playback is 10 frames/s. Scale bar=3 μm. Time is in min:s. Video corresponds to the still images from Fig. 2A.

## References

Akiyoshi B, Sarangapani KK, Powers AF, Nelson CR, Reichow SL, Arellano-Santoyo H, Gonen T, Ranish JA, Asbury CL, Biggins S. 2010. Tension directly stabilizes reconstituted kinetochore-microtubule attachments. Nature 468:576–9. doi:10.1038/nature09594

Aravamudhan P, Goldfarb AA, Joglekar AP. 2015. The kinetochore encodes a mechanical switch to disrupt spindle assembly checkpoint signalling. Nat Cell Biol 17:868–879. doi:10.1038/ncb3179

Cane S, Ye AA, Luks-Morgan SJ, Maresca TJ. 2013. Elevated polar ejection forces stabilize kinetochore-microtubule attachments. J Cell Biol 200:203–18. doi:10.1083/jcb.201211119

Cheeseman IM, Chappie JS, Wilson-Kubalek EM, Desai A. 2006. The conserved KMN network constitutes the core microtubule-binding site of the kinetochore. Cell 127:983–97. doi:10.1016/j.cell.2006.09.039

Chen RH, Shevchenko A, Mann M, Murray AW. 1998. Spindle checkpoint protein Xmad1 recruits Xmad2 to unattached kinetochores. J Cell Biol 143:283–95.

Ciferri C, Pasqualato S, Screpanti E, Varetti G, Santaguida S, Dos Reis G, Maiolica A, Polka J, De Luca JG, De Wulf P, Salek M, Rappsilber J, Moores CA, Salmon ED, Musacchio A. 2008. Implications for kinetochore-microtubule attachment from the structure of an engineered Ndc80 complex. Cell 133:427–39. doi:10.1016/j.cell.2008.03.020

Collin P, Nashchekina O, Walker R, Pines J. 2013. The spindle assembly checkpoint works like a rheostat rather than a toggle switch. Nat Cell Biol 15:1378–1385. doi:10.1038/ncb2855

De Antoni A, Pearson CG, Cimini D, Canman JC, Sala V, Nezi L, Mapelli M, Sironi L, Faretta M, Salmon ED, Musacchio A. 2005. The Mad1/Mad2 complex as a template for Mad2 activation in the spindle assembly checkpoint. Curr Biol 15:214–25. doi:10.1016/j.cub.2005.01.038

DeLuca JG, Gall WE, Ciferri C, Cimini D, Musacchio A, Salmon ED. 2006. Kinetochore microtubule dynamics and attachment stability are regulated by Hec1. Cell 127:969–82. doi:10.1016/j.cell.2006.09.047

Dick AE, Gerlich DW. 2013. Kinetic framework of spindle assembly checkpoint signalling. Nat Cell Biol 15:1370–1377. doi:10.1038/ncb2842

Drpic D, Pereira AJ, Barisic M, Maresca TJ, Maiato H. 2015. Polar Ejection Forces Promote the Conversion from Lateral to End-on Kinetochore-Microtubule Attachments on Mono-oriented Chromosomes. Cell Rep 13:460–468. doi:10.1016/j.celrep.2015.08.008

Dudka D, Noatynska A, Smith CA, Liaudet N, McAinsh AD, Meraldi P. 2018. Complete microtubule–kinetochore occupancy favours the segregation of merotelic attachments. Nat Commun 9:2042. doi:10.1038/s41467-018-04427-x

Elting MW, Hueschen CL, Udy DB, Dumont S. 2014. Force on spindle microtubule minus ends moves chromosomes. J Cell Biol 206:245–56. doi:10.1083/jcb.201401091

Elting MW, Prakash M, Udy DB, Dumont S. 2017. Mapping Load-Bearing in the Mammalian Spindle Reveals Local Kinetochore Fiber Anchorage that Provides Mechanical Isolation and Redundancy. Curr Biol 27:2112–2122.e5. doi:10.1016/j.cub.2017.06.018

Etemad B, Kuijt TEF, Kops GJPL. 2015. Kinetochore–microtubule attachment is sufficient to satisfy the human spindle assembly checkpoint. Nat Commun 6:8987. doi:10.1038/ncomms9987

Funabiki H, Wynne DJ. 2013. Making an effective switch at the kinetochore by phosphorylation and dephosphorylation. Chromosoma 122:135–58. doi:10.1007/s00412-013-0401-5

Guimaraes GJ, Dong Y, McEwen BF, Deluca JG. 2008. Kinetochore-microtubule attachment relies on the disordered N-terminal tail domain of Hec1. Curr Biol 18:1778–84. doi:10.1016/j.cub.2008.08.012

Heinrich S, Geissen E-M, Kamenz J, Trautmann S, Widmer C, Drewe P, Knop M, Radde N, Hasenauer J, Hauf S. 2013. Determinants of robustness in spindle assembly checkpoint signalling. Nat Cell Biol 15:1328–1339. doi:10.1038/ncb2864

Hengeveld RCC, Vromans MJM, Vleugel M, Hadders MA, Lens SMA. 2017. Inner centromere localization of the CPC maintains centromere cohesion and allows mitotic checkpoint silencing. Nat Commun 8:15542. doi:10.1038/ncomms15542

Hiruma Y, Sacristan C, Pachis ST, Adamopoulos A, Kuijt T, Ubbink M, von Castelmur E, Perrakis A, Kops GJPL. 2015. Competition between MPS1 and microtubules at kinetochores regulates spindle checkpoint signaling. Science (80- ) 348:1264–1267. doi:10.1126/science.aaa4055

Howell BJ, McEwen BF, Canman JC, Hoffman DB, Farrar EM, Rieder CL, Salmon ED. 2001. Cytoplasmic dynein/dynactin drives kinetochore protein transport to the spindle poles and has a role in mitotic spindle checkpoint inactivation. J Cell Biol 155:1159–72. doi:10.1083/jcb.200105093

Howell BJ, Moree B, Farrar EM, Stewart S, Fang G, Salmon ED. 2004. Spindle checkpoint protein dynamics at kinetochores in living cells. Curr Biol 14:953–64. doi:10.1016/j.cub.2004.05.053

Hueschen CL, Kenny SJ, Xu K, Dumont S. 2017. NuMA recruits dynein activity to microtubule minus-ends at mitosis. Elife 6. doi:10.7554/eLife.29328

Janssen LME, Averink T V., Blomen VA, Brummelkamp TR, Medema RH, Raaijmakers JA. 2018. Loss of Kif18A Results in Spindle Assembly Checkpoint Activation at Microtubule-Attached Kinetochores. Curr Biol 28:2685–2696.e4. doi:10.1016/J.CUB.2018.06.026

Ji Z, Gao H, Yu H. 2015. Kinetochore attachment sensed by competitive Mps1 and microtubule binding to Ndc80C. Science (80- ) 348:1260–1264. doi:10.1126/science.aaa4029

Kajtez J, Solomatina A, Novak M, Polak B, Vukušić K, Rüdiger J, Cojoc G, Milas A, Šumanovac Šestak I, Risteski P, Tavano F, Klemm AH, Roscioli E, Welburn J, Cimini D, Glunčić M, Pavin N, Tolić IM. 2016. Overlap microtubules link sister k-fibres and balance the forces on bi-oriented kinetochores. Nat Commun 7:10298. doi:10.1038/ncomms10298

Khodjakov A, Rieder CL. 1996. Kinetochores moving away from their associated pole do not exert a significant pushing force on the chromosome. J Cell Biol 135:315–27.

King JM, Nicklas RB. 2000. Tension on chromosomes increases the number of kinetochore microtubules but only within limits. J Cell Sci 113 Pt 21:3815–23.

Kuhn J, Dumont S. 2017. Spindle assembly checkpoint satisfaction occurs via end-on but not lateral attachments under tension. J Cell Biol 216:1533–1542. doi:10.1083/jcb.201611104

London N, Biggins S. 2014. Signalling dynamics in the spindle checkpoint response. Nat Rev Mol Cell Biol 15:736–748. doi:10.1038/nrm3888

Lukinavičius G, Reymond L, D’Este E, Masharina A, Göttfert F, Ta H, Güther A, Fournier M, Rizzo S, Waldmann H, Blaukopf C, Sommer C, Gerlich DW, Arndt H-D, Hell SW, Johnsson K. 2014. Fluorogenic probes for live-cell imaging of the cytoskeleton. Nat Methods 11:731–733. doi:10.1038/nmeth.2972

Magidson V, He J, Ault JG, O’Connell CB, Yang N, Tikhonenko I, McEwen BF, Sui H, Khodjakov A. 2016. Unattached kinetochores rather than intrakinetochore tension arrest mitosis in taxol-treated cells. J Cell Biol 212:307–19. doi:10.1083/jcb.201412139

Maldonado M, Kapoor TM. 2011. Constitutive Mad1 targeting to kinetochores uncouples checkpoint signalling from chromosome biorientation. Nat Cell Biol 13:475–482. doi:10.1038/ncb2223

Maresca TJ, Salmon ED. 2009. Intrakinetochore stretch is associated with changes in kinetochore phosphorylation and spindle assembly checkpoint activity. J Cell Biol 184:373–81. doi:10.1083/jcb.200808130

McCleland ML, Gardner RD, Kallio MJ, Daum JR, Gorbsky GJ, Burke DJ, Stukenberg PT. 2003. The highly conserved Ndc80 complex is required for kinetochore assembly, chromosome congression, and spindle checkpoint activity. Genes Dev 17:101–114. doi:10.1101/gad.1040903

McEwen BF, Heagle AB, Cassels GO, Buttle KF, Rieder CL. 1997. Kinetochore fiber maturation in PtK1 cells and its implications for the mechanisms of chromosome congression and anaphase onset. J Cell Biol 137:1567–80.

McKinley KL, Cheeseman IM. 2017. Large-Scale Analysis of CRISPR/Cas9 Cell-Cycle Knockouts Reveals the Diversity of p53-Dependent Responses to Cell-Cycle Defects. Dev Cell 40:405–420.e2. doi:10.1016/j.devcel.2017.01.012

Nijenhuis W, Vallardi G, Teixeira A, Kops GJPL, Saurin AT. 2014. Negative feedback at kinetochores underlies a responsive spindle checkpoint signal. Nat Cell Biol 16:1257–1264. doi:10.1038/ncb3065

O’Connell CB, Loncarek J, Hergert P, Kourtidis A, Conklin DS, Khodjakov A. 2008. The spindle assembly checkpoint is satisfied in the absence of interkinetochore tension during mitosis with unreplicated genomes. J Cell Biol 183:29–36. doi:10.1083/jcb.200801038

Orth JD, Loewer A, Lahav G, Mitchison TJ. 2012. Prolonged mitotic arrest triggers partial activation of apoptosis, resulting in DNA damage and p53 induction. Mol Biol Cell 23:567–76. doi:10.1091/mbc.E11-09-0781

Peterson JB, Ris H. 1976. Electron-microscopic study of the spindle and chromosome movement in the yeast Saccharomyces cerevisiae. J Cell Sci 22.

Pines J, Clute P. 1999. Temporal and spatial control of cyclin B1 destruction in metaphase. Nat Cell Biol 1:82–87. doi:10.1038/10049

Rieder CL, Cole RW, Khodjakov A, Sluder G. 1995. The checkpoint delaying anaphase in response to chromosome monoorientation is mediated by an inhibitory signal produced by unattached kinetochores. J Cell Biol 130:941–8.

Rieder CL, Davison EA, Jensen LC, Cassimeris L, Salmon ED. 1986. Oscillatory movements of monooriented chromosomes and their position relative to the spindle pole result from the ejection properties of the aster and half-spindle. J Cell Biol 103:581–91. doi:10.1083/JCB.103.2.581

Saurin AT, van der Waal MS, Medema RH, Lens SMA, Kops GJPL. 2011. Aurora B potentiates Mps1 activation to ensure rapid checkpoint establishment at the onset of mitosis. Nat Commun 2:316. doi:10.1038/ncomms1319

Shrestha RL, Conti D, Tamura N, Braun D, Ramalingam RA, Cieslinski K, Ries J, Draviam VM. 2017. Aurora-B kinase pathway controls the lateral to end-on conversion of kinetochore-microtubule attachments in human cells. Nat Commun 8:150. doi:10.1038/s41467-017-00209-z

Sikirzhytski V, Magidson V, Steinman JB, He J, Le Berre M, Tikhonenko I, Ault JG, McEwen BF, Chen JK, Sui H, Piel M, Kapoor TM, Khodjakov A. 2014. Direct kinetochore-spindle pole connections are not required for chromosome segregation. J Cell Biol 206:231–43. doi:10.1083/jcb.201401090

Stumpff J, Wagenbach M, Franck A, Asbury CL, Wordeman L. 2012. Kif18A and chromokinesins confine centromere movements via microtubule growth suppression and spatial control of kinetochore tension. Dev Cell 22:1017–29. doi:10.1016/j.devcel.2012.02.013

Suzuki A, Badger BL, Salmon ED. 2015. A quantitative description of Ndc80 complex linkage to human kinetochores. Nat Commun 6:8161. doi:10.1038/ncomms9161

Tauchman EC, Boehm FJ, DeLuca JG. 2015. Stable kinetochore–microtubule attachment is sufficient to silence the spindle assembly checkpoint in human cells. Nat Commun 6:10036. doi:10.1038/ncomms10036

Uchida KSK, Takagaki K, Kumada K, Hirayama Y, Noda T, Hirota T. 2009. Kinetochore stretching inactivates the spindle assembly checkpoint. J Cell Biol 184:383–90. doi:10.1083/jcb.200811028

Uetake Y, Sluder G. 2010. Prolonged prometaphase blocks daughter cell proliferation despite normal completion of mitosis. Curr Biol 20:1666–71. doi:10.1016/j.cub.2010.08.018

Waters JC, Chen RH, Murray AW, Salmon ED. 1998. Localization of Mad2 to kinetochores depends on microtubule attachment, not tension. J Cell Biol 141:1181–91. doi:10.1083/JCB.141.5.1181

Wendell KL, Wilson L, Jordan MA. 1993. Mitotic block in HeLa cells by vinblastine: ultrastructural changes in kinetochore-microtubule attachment and in centrosomes. J Cell Sci 104 (Pt 2):261–74.

Yoo TY, Choi J-M, Conway W, Yu C-H, Pappu R V, Needleman DJ. 2018. Measuring NDC80 binding reveals the molecular basis of tension-dependent kinetochore-microtubule attachments. Elife 7. doi:10.7554/eLife.36392

Zaytsev A V, Mick JE, Maslennikov E, Nikashin B, DeLuca JG, Grishchuk EL. 2015. Multisite phosphorylation of the NDC80 complex gradually tunes its microtubule-binding affinity. Mol Biol Cell 26:1829–44. doi:10.1091/mbc.E14-11-1539

Zaytsev A V, Sundin LJR, DeLuca KF, Grishchuk EL, DeLuca JG. 2014. Accurate phosphoregulation of kinetochore-microtubule affinity requires unconstrained molecular interactions. J Cell Biol 206:45–59. doi:10.1083/jcb.201312107

